# Retrospective cues mitigate information loss in human cortex during working memory storage

**DOI:** 10.1101/351544

**Authors:** Edward F. Ester, Asal Nouri, Laura Rodriguez

## Abstract

Working memory (WM) enables the flexible representation of information over short intervals. It is well-established that WM performance can be enhanced by a retrospective cue presented during storage, yet the neural mechanisms responsible for this benefit are unclear. Here, we tested several explanations for retro-cue benefits by quantifying changes in spatial WM representations reconstructed from alpha-band (8-12 Hz) EEG activity recorded from human participants (both sexes) before and after presentation of a retrospective cue. This allowed us to track cue-related changes in WM representations with high temporal resolution (tens of milliseconds). Participants encoded the locations of two colored discs for subsequent report. During neutral trials an uninformative cue instructed participants to remember the locations of both discs across a blank delay, and we observed a monotonic decrease in the fidelity of reconstructed spatial WM representations with time. During valid trials a 100% reliable cue indicated the color of the disc participants would be probed to report. Critically, valid cues were presented immediately after termination of the encoding display (“valid early”, or VE trials) or midway through the delay period (“valid late” or VL trials). During VE trials the gradual loss of location-specific information observed during neutral trials was eliminated, while during VL trials it was partially reversed. Our findings suggest that retro-cues engage several different mechanisms that together serve to mitigate information loss during WM storage.

**Significance Statement:** Working memory (WM) performance can be improved by a cue presented during storage. This effect, termed a retrospective cue benefit, has been used to explore the limitations of attentional prioritization in WM. However, the mechanisms responsible for retrospective cue benefits are unclear. Here we tested several explanations for retrospective cue benefits by examining how they influence WM representations reconstructed from human EEG activity. This approach allowed us to visualize, quantify, and track the effects of retrospective cues with high temporal resolution (on the order of tens of milliseconds). We show that under different circumstances retrospective cues can both eliminate and even partially reverse information loss during WM storage, suggesting that retrospective cue benefits have manifold origins.

---

Visual working memory (WM) enables the representation and manipulation of information no longer in the sensorium. This system is an integral component of many higher-order cognitive abilities (e.g., Cowan et al. 2000), yet it has a sharply limited representational capacity (e.g., Luck & Vogel, 2013; Ma et al. 2014). Thus, mechanisms of selective attention are needed to control access to WM (e.g., Vogel et al. 2005) and to prioritize existing WM representations for behavioral output (e.g., Myers et al. 2017). Attentional prioritization in WM has been extensively studied using retrospective cues (see Souza & Oberauer, 2016 and Myers et al. 2017 for recent comprehensive reviews). In a typical retro-cue experiment, participants encode an array of items for subsequent report. During the ensuing delay period a cue indicates which of the original items is most likely to be tested. Relative to a no-cue or neutral-cue baseline, valid cues typically yield greater memory performance while invalid cues typically yield reduced memory performance (though the evidence for invalid cue costs is mixed; see Souza & Oberauer 2016).

Several mechanisms may be responsible for retrospective cue benefits in WM performance. For example, multiple studies have reported reductions in load-dependent neural signatures of WM storage following a retrospective cue, suggesting that these cues engage mechanisms that facilitate the removal of irrelevant items from WM (e.g., Kuo et al. 2012; Williams & Woodman, 2012). Other studies have reported changes in lateralized alpha band activity when participants are retrospectively cued to an item that previously appeared in the left or right visual hemifields, consistent with an attentional prioritization of the cued representation (and/or suppression of the uncued representation; e.g., Poch et al. 2014; Myers et al. 2015; van Moorselaar et al. 2018). There is also evidence suggesting that valid retrospective cues trigger mechanisms that insulate cued WM representations from subsequent display or interference (e.g., Makovski et al., 2010; Pertzov et al. 2013), or mechanisms that facilitate comparisons between cued WM representations and subsequent memory probes (e.g., Souza & Oberauer 2016).

Here, we tested several explanations for retrocue benefits by quantifying changes in spatially-specific WM representations before and after the appearance of a retrospective cue. Inspired by earlier work (e.g., Foster et al. 2016), we reconstructed spatially-specific mnemonic representations by applying an inverted encoding model (IEM) to spatiotemporal patterns of alpha band (8-12 Hz) activity recorded while participants performed a retrospectively cued spatial WM task. On each trial participants encoded the locations of two colored discs (blue and red). During neutral trials, an uninformative color cue presented after the encoding display informed participants to remember the locations of both discs across a blank interval. During valid trials a 100% reliable color cue indicated which disc would be probed at the end of the trial. Valid cues were presented either immediately after offset of the encoding display (“Valid Early” trials, VE), or at the midpoint of the subsequent blank period (“Valid Late” trials; VL). We isolated the effects of retrospective cues on spatial WM performance by comparing location-specific WM representations across the neutral and valid conditions. To preview the results, during neutral trials we observed a gradual decrease in the fidelity of location-specific representations with time. During VE trials this decrease was eliminated, while during VL trials it was partially reversed. Our findings thus support the view that retro-cues can engage several different mechanisms that together serve to mitigate information loss during WM storage.

## Methods

### Participants

31 volunteers from the Florida Atlantic University community (ages 18-40, both sexes) completed a single 2.5-hour testing session. All participants self-reported normal or corrected-to-normal visual acuity and were compensated at a rate of $15/hr. Data from 4 participants were discarded due to an excessive number of electrooculogram artifacts (over 33% of trials). Data from a fifth participant was discarded as s/he withdrew from the study after completing only six of twelve testing blocks. The data reported here reflect the remaining 26 volunteers.

### Testing Environment

Participants were seated in a dimly-lit and sound-attenuated (unshielded) recording chamber. Stimuli were generated in MATLAB using Psychtoolbox-3 software (Kleiner et al. 2007) and rendered on a 17-inch Dell CRT monitor cycling at 85 Hz. Participants were seated approximately 60 cm from the display (head position was not constrained).

### Spatial Memory Task

The behavioral task was based on an experimental approach described by Sprague et al. (2016). A trial schematic is shown in Figure 1. Participants were instructed to fixate a small dot (subtending 0.2° from a viewing distance of 60 cm) throughout the experiment. Each trial began with a sample display containing two colored discs (one red and one blue). Each disc was presented in one of 9 equally spaced positions (40° to 360° in 40° increments) along the perimeter of an imaginary circle (radius 6° visual angle) centered at the fixation point. A small amount of jitter (±10° polar angle) was added to the location of each disc on each trial to discourage verbal coding strategies (e.g., “the blue disc was at 2 o’clock”).

**Figure 1.**
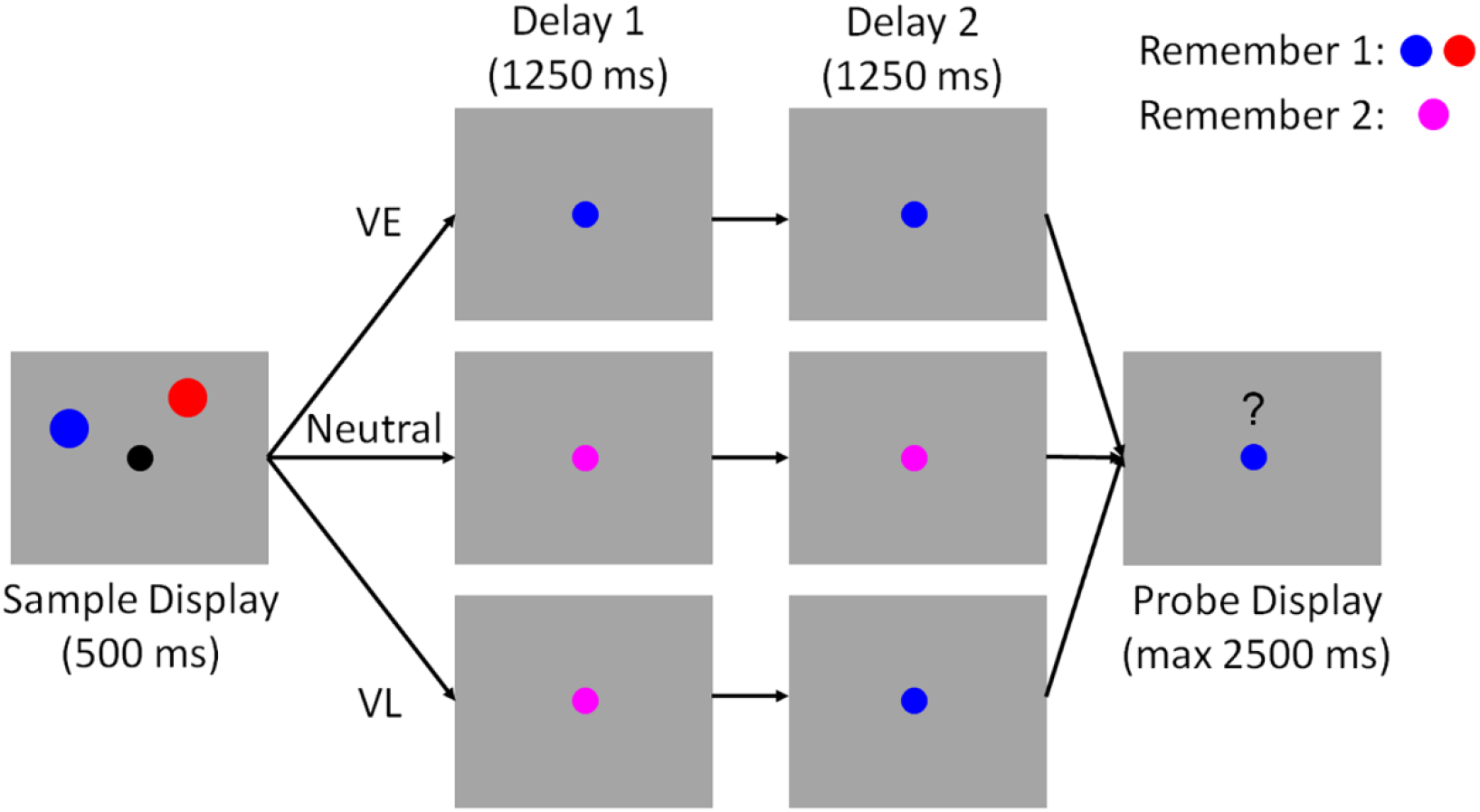
Spatial Memory Task. Participants encoded the locations of two colored discs (red and blue; sample display). Each disc was presented at one of nine equally spaced locations along the perimeter of an imaginary circle centered at fixation (see Methods and Figure 3). During valid-early trials (VE), the color of the fixation point changed from black to either blue or red immediately after termination of the sample display. This cue indicated which disc would be probed with 100% reliability. During neutral trials, the color of the fixation point changed from black to purple, instructing participants to remember the locations of both discs. During valid-late (VL) trials, the color of the fixation point changed from black to purple. Halfway through the subsequent delay period, the fixation point changed colors from purple to either blue or red. This second change indicated which disc would be probed with 100% reliability. Each trial concluded with a probe display containing a red or blue fixation point, a question mark, and a mouse cursor. Participants were instructed to report the precise location of the disc indicated by the color of the fixation point via mouse click. Note: the above schematic is included for illustrative purposes; displays are not drawn to scale. See *Methods* for display and stimulus geometry.

The sample display was extinguished after 500 ms. During “Valid Early” (VE) trials the fixation point changed colors from black to either blue or red. This change served as a 100% valid cue for the disc whose location participants would be asked to report at the end of the trial. The fixation point remained blue (or red) for the remainder of the delay period. During “Valid Late” trials (VL), the color of the fixation point initially changed from black to purple, instructing participants to remember the locations of both discs. At the midpoint of the delay period (1250 ms after offset of the sample display) the fixation point changed colors from purple to either blue or red; this change served as a 100% valid cue for the disc whose location participants would be asked to report at the end of the trial. The fixation dot remained red or blue for the remainder of the second delay period (1251-2500 ms after sample offset). Finally, during neutral trials the color of the fixation point changed from black to purple and remained purple across the entire 2500 ms delay period.

Each trial concluded with a test display containing a blue or red fixation point, a mouse cursor, and a question mark symbol (“?”) above the fixation point. Participants were required to click on the location of the disc indicated by the color of the fixation point within a 3000 ms response window. Memory performance was quantified as the absolute angular distance between the polar location of the probed disc and the polar location reported by the participant. Performance feedback (mean absolute recall error; i.e., the mean absolute difference between the polar angle reported by the participant and the polar location of the probed disc) was given at the end of each block. Participants completed 10 (N = 1), 11 (N = 2), or 12 (N = 23) blocks of 72 trials. Cue conditions (VE, VL, neutral) were counterbalanced within each block, while the spatial positions of the red and blue discs were counterbalanced across sub-sessions of six blocks.

### EEG Acquisition and Preprocessing

Continuous EEG was recorded from 62 scalp electrodes via a BrainProducts actiCHamp amplifier (Munich, Germany). Additional electrodes were placed over the left and right mastoids. Electrode impedances were kept below 15 kΩ. Data were recorded with a right mastoid reference and digitized at 1000 Hz. The horizontal electrooculogram (EOG) was recorded from electrodes located ~1 cm from the left and right canthi while the vertical EOG was recorded from electrodes placed above and below the right eye. Data were later re-referenced to the algebraic mean of the left and right mastoids, bandpass filtered from 0.1 to 40 Hz (3^rd^ order zero-phase forward-and-reverse Butterworth filters), epoched from −1000 to +4000 ms relative to the start of each trial, and baseline-corrected from - 100 to 0 ms relative to the start of each trial. Trials contaminated by blinks or horizontal eye movements greater than ~2.5° (assuming a normative voltage threshold of 16 μV/°; Lins et al. 1993) were excluded from subsequent analyses. Noisy electrodes were identified and removed by visual inspection. Across participants, we rejected an average (±1 S.E.M.) of 12.38% (±1.83%) trials and 1.04 (±0.23) electrodes.

### Inverted Encoding Model

Following earlier work (e.g., Foster et al. 2016) we used an inverted encoding model to reconstruct location-specific representations of the red and blue discs during the sample display and subsequent delay period. Reconstructions of stimulus locations were computed from the spatial topography of induced alpha-band (8-12 Hz) power measured across 17 occipitoparietal electrode sites: O1, O2, Oz, PO7, PO3, POz, PO4, PO8, P7, P5, P3, P1, Pz, P2, P4, P6, and P8. Robust representations of spatial position could also be reconstructed from frontal, central, and temporal electrode sites, but these representations were substantially weaker than those reconstructed from occipitoparietal electrode sites. To isolate alpha-band activity, the raw EEG time series at each electrode was bandpass filtered from 8-12 Hz (3^rd^ order zero-phase forward-and-reverse Butterworth), yielding a real-valued signal *f*(*t*). The analytic representation of *f*(*t*) was obtained by applying a Hilbert transformation:

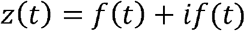

where *i* = √-1 and *if*(*t*) = *A*(*t*)*e*^*iφ*(*t*)^. Induced alpha power was computed by extracting and squaring the instantaneous amplitude *A*(*t*) of the analytic signal *z*(*t*).

We modeled alpha power at each scalp electrode as a weighted sum of 9 location-selective channels, each with an idealized tuning curve (a half-wave rectified cosine raised to the 8^th^ power). The maximum response of each channel was normalized to 1, thus units of response are arbitrary. The predicted responses of each channel during each trial were arranged in a *k* channel by *n* trials design matrix *C*. Separate design matrices were constructed to track the locations of the blue and red discs across trial (i.e., we reconstructed the locations of the blue and red discs separately, then later sorted these reconstructions according to cue condition).

The relationship between the data and the predicted channel responses *C* is given by a general linear model of the form:

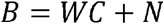

where B is a *m* electrode by *n* trials training data matrix, W is an *m* electrode by *k* channel weight matrix, and *N* is a matrix of residuals (i.e., noise).

To estimate *W*, we constructed a “training” data set containing an equal number of trials from each stimulus location (i.e., 40-360° in 40° steps). We first identified the location *φ* with the fewest *r* repetitions in the full data set after EOG artifact removal. Next, we constructed a training data set *B_trn_* (*m* electrodes by *n* trials) and weight matrix *C_trn_* (*n* trials by *k* channels) by randomly selecting (without replacement) 1:*r* trials for each of the nine possible stimulus locations (ignoring cue condition; i.e., the training data set contained a mixture of VE, VL, and Neutral trials). The training data set was used to compute a weight for each channel *C_i_* via least-squares estimation:

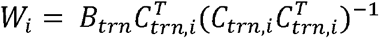

where *C_m,i_* is an *n* trial row vector containing the predicted responses of spatial channel *i* during each training trial.

The weights *W* were used to estimate a set of spatial filters *V* that capture the underlying channel responses while accounting for correlated variability between electrode sites (i.e., the noise covariance; Kok et al. 2017):

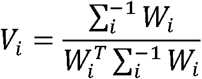

where *Σ_i_* is the regularized noise covariance matrix for channel *i* and estimated as:

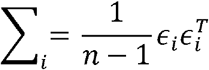

where *n* is the number of training trials and ***ε**_i_* is a matrix of residuals:

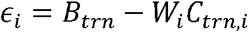

Estimates of *ε_i_* were obtained by regularization-based shrinkage using an analytically determined shrinkage parameter (see Blankertz et al. 2011; Kok et al. 2017). An optimal spatial filter *v_i_* was estimated for each channel *C_i_*, yielding an *m* electrode by *k* filter matrix *V*.

Next, we constructed a “test” data set *B_tst_* (*m* electrodes by *n* trials) containing data from all trials not included in the training data set and estimated trial-by-trial channel responses *C_tst_* (*k* channels x *n* trials) from the filter matrix *V* and the test data set:

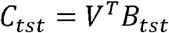

Trial-by-trial channel responses were interpolated to 360°, circularly shifted to a common center (0°, by convention), and sorted by cue condition (i.e., VE, VL, neutral). To quantify the effects of retrospective cues on spatial memory representations, we obtained an estimate of location specific information by converting the (averaged) channel response estimates for each cue condition to polar form and projected them onto a vector with angle 0°:

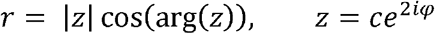

where *c* is a vector of estimated channel responses and *φ* is the vector of angles at which the channels peak.

To ensure internal reliability this entire analysis was repeated 100 times, and unique (randomly chosen) subsets of trials were used to define the training and test data sets during each permutation. The results were then averaged across permutations.

### Estimating the Temporal Resolution of Location Information Estimates

As described in the preceding section, location-specific reconstructions (and hence, estimates of location information) were computed using instantaneous distributions of alpha power across electrode sites at each time point. However, the bandpass filter used to isolate alpha band activity (8-12 Hz) can introduce temporal distortions, making it difficult to make precise statements about the temporal resolution of the analysis pipeline. To formally investigate this issue, we generated a 3000 ms sinusoid with unit amplitude and a frequency of 10 Hz (along with 1000 ms of pre- and post-signal zero padding for a total signal length of 5000 ms) and ran it through the filtering routine used to identify alpha-band activity (defined as 8-12 Hz) in the raw EEG signal. Temporal distortions caused by filtering were defined as points where the amplitude of the filtered signal reached 25% of maximum (1.0). For a perfect filter, these points would occur at precisely 1000 and 4000 ms relative to stimulus onset. In reality, we obtained estimates of 963 4039 ms, or 37 ms prior to and 39 ms after stimulus onset. Thus, we estimated the temporal resolution of our analysis path as approximately ±40 ms. That is, if estimates of location information are significantly above zero at a given time point, then we can be certain that the spatial distribution of alpha power across electrode sites contained robust information about that location at some point within ±40 ms.

### Statistical Comparisons

We used a nonparametric sign permutation test to quantify differences between estimates of location information across stimuli (cued vs. uncued) and cue conditions (i.e., VE, VL, and neutral trials). Each test we performed – e.g., whether estimated location information is reliably above zero or whether estimated location information is higher for the cued vs. uncued disc – assumes a null statistic of 0. We therefore generated a null distribution of location information estimates by randomly inverting the sign of each participant’s data (with 50% probability) and averaging the data across participants. This procedure was repeated 10,000 times, yielding a 10,000-element null distribution for each time point. Finally, we implemented a cluster-based permutation test with cluster-forming and cluster-size thresholds of *p* < 0.05 to correct for multiple comparisons across time points (see Maris & Oostenveld, 2011; Wolff et al. 2017).

### Eye Movement Control Analyses

Although we excluded trials contaminated by horizontal eye movements, small but reliable biases in eye position towards the location(s) of the remembered disc(s) could nevertheless contribute to reconstructions of stimulus location. We examined this possibility in two complementary analyses. In the first analysis, we computed angular measurements of eye position by converting trial-by-trial horizontal and vertical EOG recordings to polar format (using normative scaling values of 16 μV/° and 12 μV/° for the horizontal and vertical EOG channels, respectively; Lins et al. 1993; Bae & Luck 2018), then constructed circular histograms of angular estimates of eye position as a function of stimulus location. Angular eye position estimates at each timepoint were sorted into nine bins whose centers matched the possible stimulus positions (i.e., 40° to 360° in 40° increments). For simplicity we restricted our analysis to VE trials where consistent biases in eye position should be most apparent (i.e., because participants were only required to remember one location).

In the second analysis, we regressed trial-by-trial horizontal EOG recordings (in μV; see Foster et al. 2016) onto the horizontal position of the remembered disc, again focusing on VE trials. Positive regression coefficients thus reflect greater changes in eye position as a function of stimulus location. Separate regressions were run for each participant and the resulting coefficients were averaged across participants.

### Quantifying Sources of Location Information Loss and Recovery

We evaluated potential sources of change in location-specific representations via curve-fitting analyses (e.g., Ester et al. 2015; Ester et al. 2016). We first computed a one-dimensional reconstructed representation of location for each participant by averaging channel responses over time (separately for the first and second delay periods). Reconstructions were averaged across participants and fit with a circular function of the form

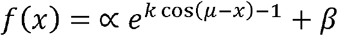

where *x* is a vector of polar locations and *β* is an additive scaling parameter. The additional parameters α, μ, and k control the amplitude, center, and concentration (i.e., the inverse of bandwidth) of the function. α, β, μ, and *k* were estimated separately for each condition via an iterative minimization algorithm (Nelder-Mead simplex as implemented by MATLAB’s “fminsearch” function).

Differences in parameters across conditions (i.e., first vs. second delay and neutral vs. VE vs. VL) were evaluated using a bootstrap test. For each comparison we selected (with replacement) and averaged 26 of 26 participant reconstructions from the two conditions of interest (e.g., neutral delay 1 and VE delay 1). The resulting average functions were fit with the circular function described above, yielding a set of parameter estimates for each condition. This procedure was repeated 10,000 times, and we computed empirical p-values for differences in parameter estimates across conditions by estimating the total proportion of permutations where parameter estimates in one condition were greater than (or less than) estimates for the other condition.

### Within-participant Variability

We report estimates of within-participant variability (e.g., 95% within-participant confidence intervals) throughout the paper. These estimates discard subject variance (e.g., overall differences in response strength) and instead reflect variance related to the subject by condition(s) interaction term(s) (e.g., variability in response strength across experimental conditions; Loftus & Masson, 1994; Cousineau 2005). We used the approach described by Cousineau (2005): raw data (e.g., location information or channel response estimates) were de-meaned on a participant by participant basis, and the grand mean across participants was added to each participant’s zero-centered data. The grand mean centered data were then used to compute bootstrapped within-participant confidence intervals (10,000 permutations).

## Results

### Valid Retrocues Enhance Memory Performance

We recorded EEG while participants performed a retrospectively cued spatial WM task (Figure 1). During neutral trials, an uninformative cue instructed participants to remember the locations of two colored discs across a blank delay period. During valid trials a 100% reliable color cue indicated which disc would be probed at the end of the trial. Valid cues were presented immediately after termination of the encoding display (valid early trials; VE) or midway through the subsequent blank delay period (valid late trials; VL). At the end of the trial, participants recalled the location of the probed disc via a mouse click. Behavioral performance was quantified as mean absolute recall error (i.e., the mean absolute difference between the polar angle reported by the participant and the polar location of the probed disc). Figure 2 plots behavioral performance as a function of trial type (i.e., neutral, VE, VL). Consistent with earlier findings (Sprague et al., 2016), average absolute recall error was reliably lower during VE and VL trials (M = 7.75° and 8.26°, respectively) relative to neutral trials (M = 8.96°; false-discovery rate corrected bootstrap tests, *p* < 0.001). Average absolute recall error was also reliably lower during VE relative to VL trials (bootstrap test; *p* < 0.001). Thus, valid retro-cues improved WM performance.

**Figure 2.**
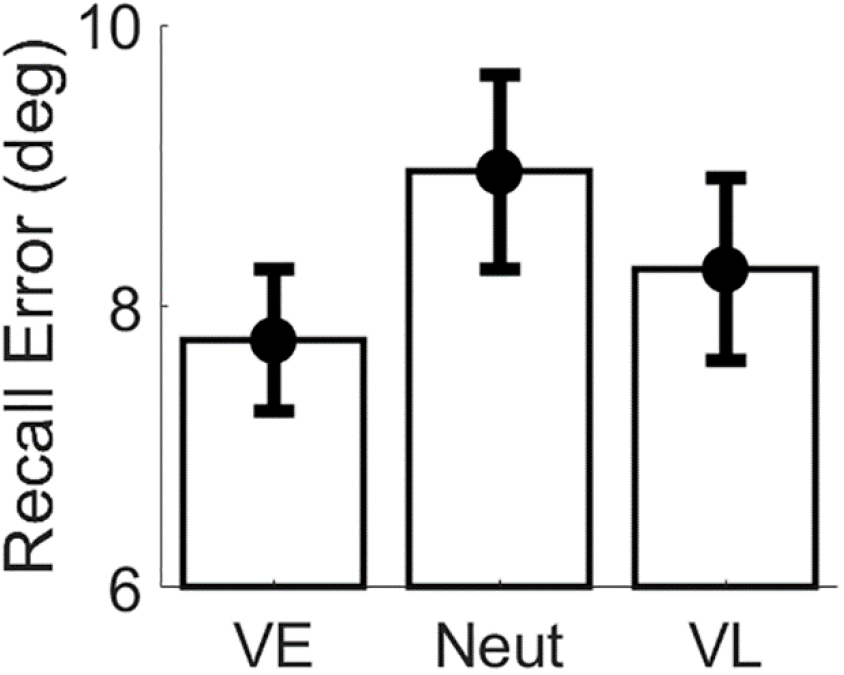
Behavioral Performance. Recall error (i.e., the absolute polar angle between the location reported by the participant and the location of the probed disc) was reliably lower during VE relative to Neutral and VL trials. Recall error was also reliably lower during VL relative to Neutral trials. Error bars show 95% confidence intervals.

### Reconstructing Location-specific WM Representations from Alpha-band Activity

Building upon earlier work (Foster et al. 2016; Samaha et al. 2016), we used an inverted encoding model to reconstruct the locations of the cued and uncued discs from spatiotemporal patterns of induced α-band power measured across occipitoparietal electrode sites (see *Inverted Encoding Model*, Methods). Briefly, we modeled instantaneous induced α-power at each electrode site as a weighted combination of 9 location-selective channels, each with an idealized response function.

The resulting channel weights were then used to calculate a predicted response for each channel given spatiotemporal distributions of α-power measured during an independent test data set. Separate reconstructions were computed for each disc irrespective of cue condition, and the individual reconstructions were averaged to yield a single time-resolved representation of location-specific activity. Consistent with earlier findings (Foster et al. 2016) trial-by-trial variability in channel responses reliably tracked the angular locations of the cued and uncued discs (Figure 3A). For convenience, we circularly shifted reconstructions for each stimulus location to a common center (0° by convention) and averaged the centered reconstructions across locations, yielding a single time-resolved reconstruction. Finally, we converted these centered reconstructions to polar form and projected them onto a unit vector with an angle of 0°. As shown in Figure 3C, location information increased rapidly following presentation of the sample display and reached an asymptotic limit approximately 400-500 ms later. During the subsequent delay period location information gradually decreased with time, though overall information levels remained reliably above 0 for the duration of the trial. Thus, our analytical approach allowed us to visualize spatially-specific WM representations with high temporal precision.

**Figure 3.**
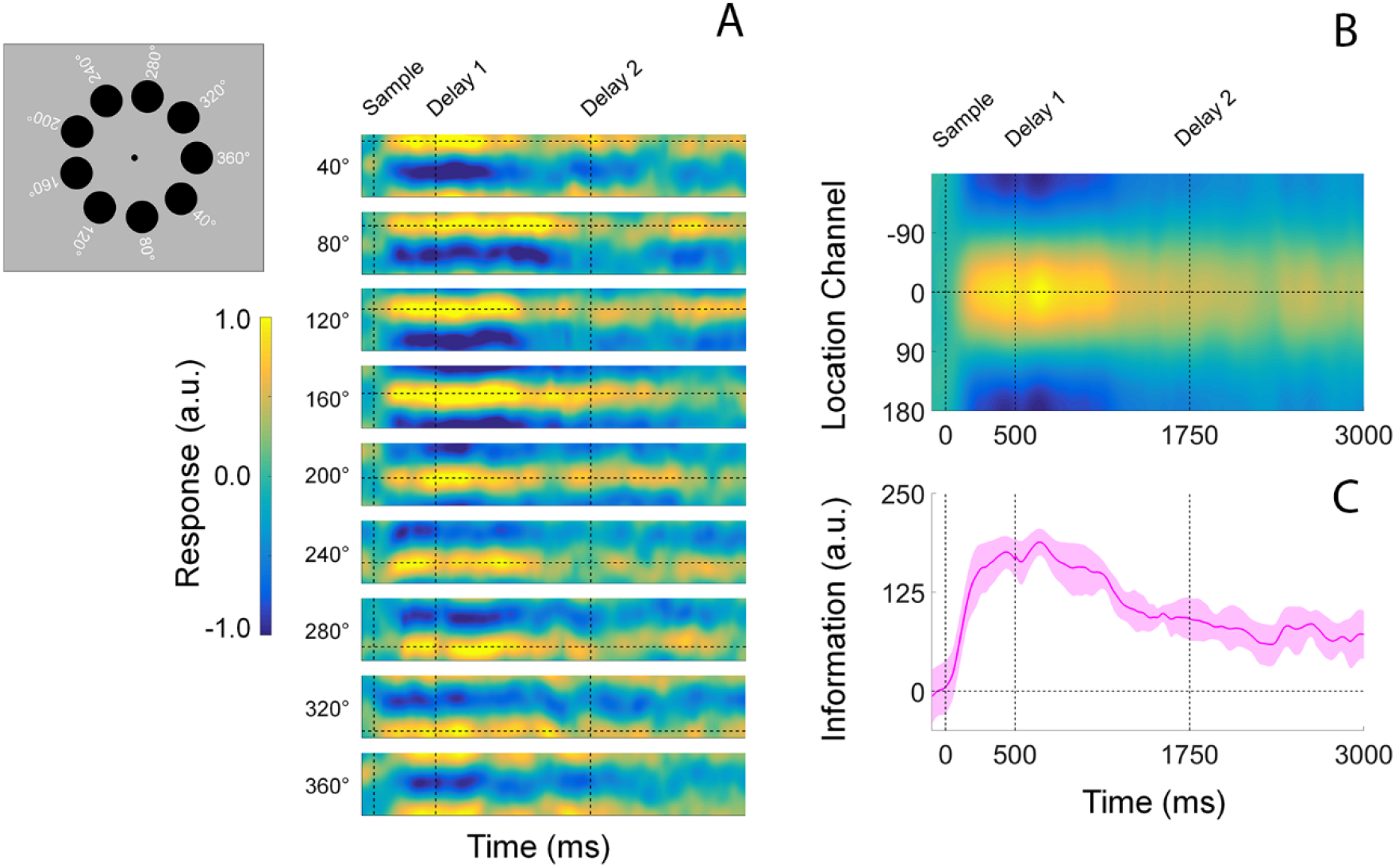
Computing time-resolved reconstructions of stimulus location. (A) Location reconstructions for each of the 9 possible stimulus locations from 40°-360°. The dashed horizontal line in each plot shows the polar location of the stimulus, while the vertical dashed lines at time 0, 500, and 1750 mark the start of the sample epoch, first delay period, and second delay period, respectively (time axis is identical to that shown in Panels B and C). The inset to the left shows each of the nine possible disc locations on each trial. Reconstructions have been pooled and averaged across stimulus identity (i.e., red vs. blue disc) and cue condition (VE, VL, neutral). (B) We circularly shifted the reconstructions shown in (A) to a common center (0°) and averaged them, yielding a single time-resolved location reconstruction. Response scale is identical to that shown in (A). (C) We converted the reconstructions shown in (B) to polar form and projected them onto a vector with angle 0°. We interpreted the resultant vector length as a measure of total location-specific information. Shaded regions represent the 95% within-participant confidence interval. a.u., arbitrary units.

### Comparison of Spatial Representations During Encoding

The central goal of this study was to examine the effects of retrospective cues on reconstructed spatial WM representations. Although participants had no way of knowing what type of cue would be presented on each trial, it is conceivable that differences in reconstructions across cue conditions during the encoding phase of the trial contributed to subsequent differences across retrospective cue conditions. We evaluated this possibility by directly comparing estimates of location information across retrospective cue conditions (VE, VL, and Neutral; Figure 4A). As shown in Figure 4B, estimates of location information were remarkably consistent across cue conditions, and we failed to identify any statistically robust differences in location information as a function of cue condition over the entire 500 ms encoding period (false-discovery-rate-corrected cluster-based permutation tests, p > 0.05). Thus, we can be certain that any effects of cue type during the subsequent WM period are not due to differences that emerged during encoding.

**Figure 4.**
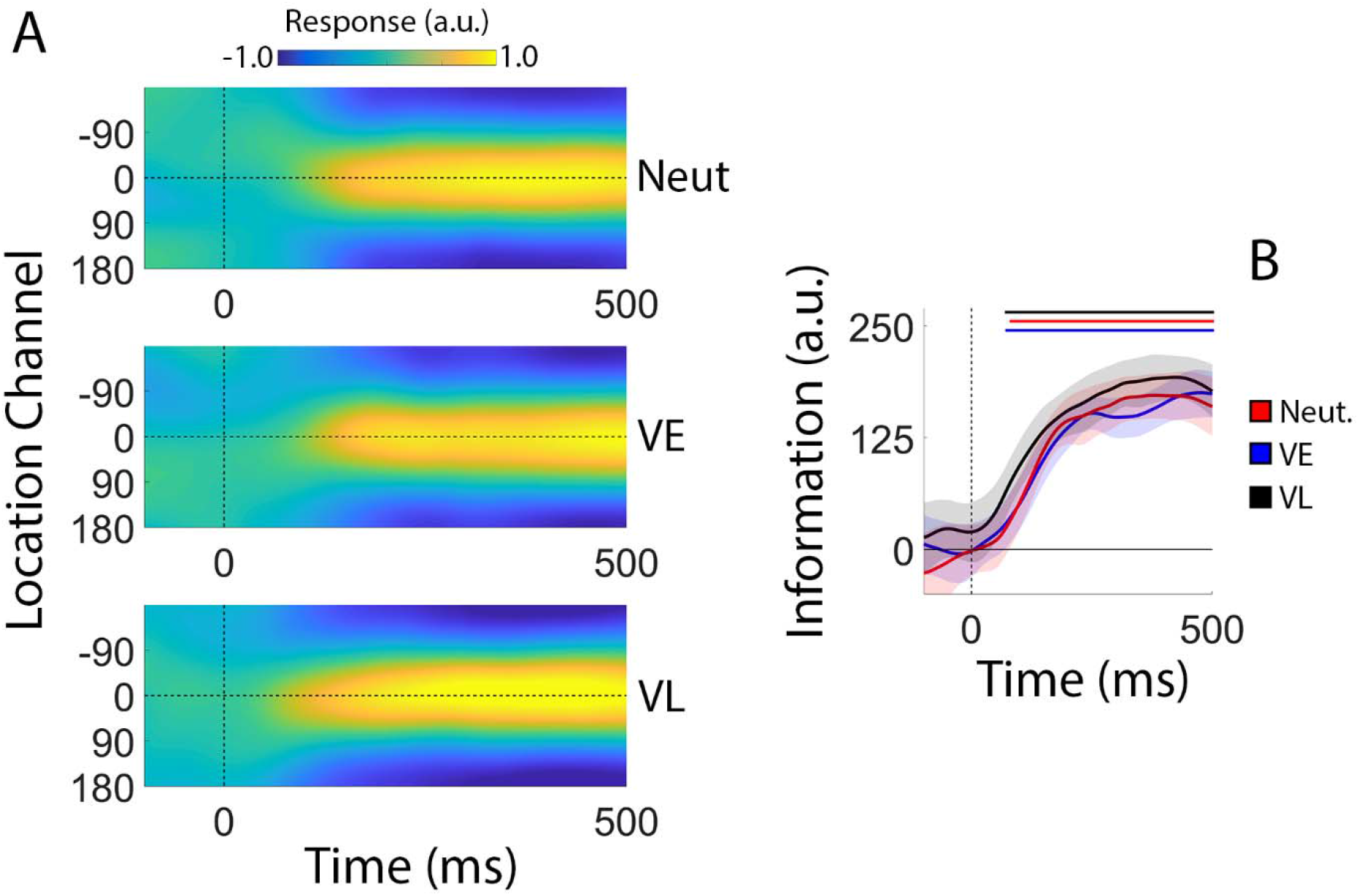
Stimulus Reconstructions and Location Information during Encoding. Participants encoded the locations of two colored discs for subsequent report (see Figure 1). (A) Channel response functions were identical across cue conditions. This is unsurprising, as participants had no way of knowing what type of cue they would receive until termination of the encoding display. (B) Robust location information emerged approximately 100 ms after display onset and increased rapidly before reaching an asymptotic limit approximately 350 ms after display onset. Statistically, estimates of location information were identical across cue conditions. Thus, we can be certain that differences in location information during the subsequent WM period are not due to differences that emerge during encoding. Solid lines at the top of panel B mark epochs where estimates of location information were reliably greater than 0. Shaded regions depict the 95% within-participant confidence interval of the mean. a.u., arbitrary units.

### Degradation of Spatial WM Representations During Neutral Trials

Next, we examined the effects of retro-cues cues on reconstructed spatially-specific WM representations. Since all retrospective cues were presented after the offset of the sample display, we limited our analyses to the 2500 ms blank interval separating the offset of the sample display and the onset of the probe display.

During neutral trials, a retrospective cue presented immediately after offset of the sample display instructed participants to remember the locations of both discs. As shown in Figure 5A, location information decreased monotonically over the course of the delay period (linear slope = −62.97 units/sec; *p* < 0.002, bootstrap test) with information reaching levels indistinguishable from 0 by the onset of the probe display. We next examined possible sources of information loss, including reductions in reconstruction amplitude (i.e., a lower signal-to-noise ratio; Sprague et al. 2014; 2016) or increases in reconstruction bandwidth (i.e., a loss of spatial precision; Ester et al. 2013). We averaged each participant’s time-resolved location reconstructions over periods from 0-1250 ms and 1251-2500 ms after the offset of the sample display (Figure 5B). Each reconstruction was fit with a circular function containing free parameters for amplitude (i.e., maximum over baseline response), concentration (the inverse of bandwidth), and center (see *Quantifying Sources of Information Loss and Recovery*, Methods). As shown in Figure 5C, we observed a decrease in response amplitude across the first and second half of the delay period, but no change in concentration. Thus, requiring participants to store multiple spatial WM representations was associated with a gradual decrease in the strength of each representation over time.

**Figure 5.**
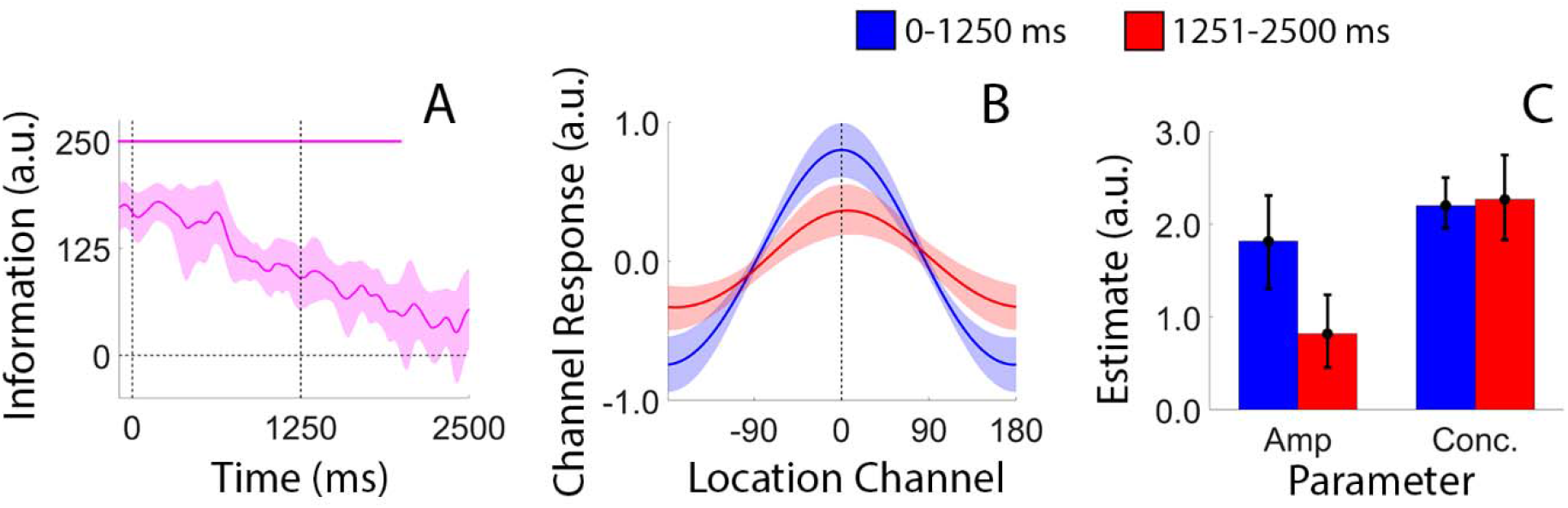
Degradation of Spatial WM Representations During Neutral Trials. (A) During neutral trials we observed a monotonic decrease in total location information during the delay period. Data have been pooled and averaged across stimulus locations (i.e., the locations of the blue and red discs) and are time-locked to the offset of the sample display (0 ms). The pink bar at the top of the plot marks epochs where estimates of location information were reliably greater than zero (false-discovery-rate-corrected cluster-based permutation test, see Methods). (B) We identified source(s) of information loss by computing and quantifying time-averaged location-specific reconstructions during the first and second delay periods (see *Quantifying Sources of Information Loss and Recovery*, Methods). (C) Reconstruction amplitudes were reliably lower during the second relative to the first delay period, suggesting that information loss reflects a gradual reduction in the overall strength of each spatial WM representation. For all plots shaded regions and error bars show 95% within-participant confidence intervals. a.u., arbitrary units.

### Valid Retrocues Presented Immediately After Encoding Eliminate Information Loss

During VE trials, a retrospective cue presented immediately after the sample display indicated which disc (blue or red) would be probed with 100% certainty. In contrast to the pattern seen during neutral trials (Figure 5), location information for the cued disc remained constant across the delay period (Figure 6; linear slope = −17.8 units/sec; *p* = 0.06; bootstrap test). Contrary to prior results suggesting a cue-driven increase in information over-and-above that observed during encoding (e.g., Rerko et al., 2014; Schneegans & Bays, 2017), we found no evidence for a strengthening of the cued location immediately after cue onset (obtained by comparing average location information over epochs spanning the last 100 ms of the sample period and the first 500 ms of the delay period; *p* = 0.93, bootstrap test). Conversely, location information for the uncued disc quickly fell to 0. The rate of information loss for the uncued disc during the first 1250 ms of the delay period during VE trials was nearly double that observed for during the same interval of neutral trials (linear slopes of −143.61 vs. 74.18 units/sec; *p* < 0.02; bootstrap test). Clear differences between location information estimates for the cued and uncued discs emerged approximately 600 ms after cue onset and persisted throughout the remainder of the delay period (Figure 6B; red markers). This delay is consistent with behavioral studies suggesting that it takes participants approximately 300-600 ms to process and utilize a retrospective cue (e.g., Souza et al., 2014).

**Figure 6.**
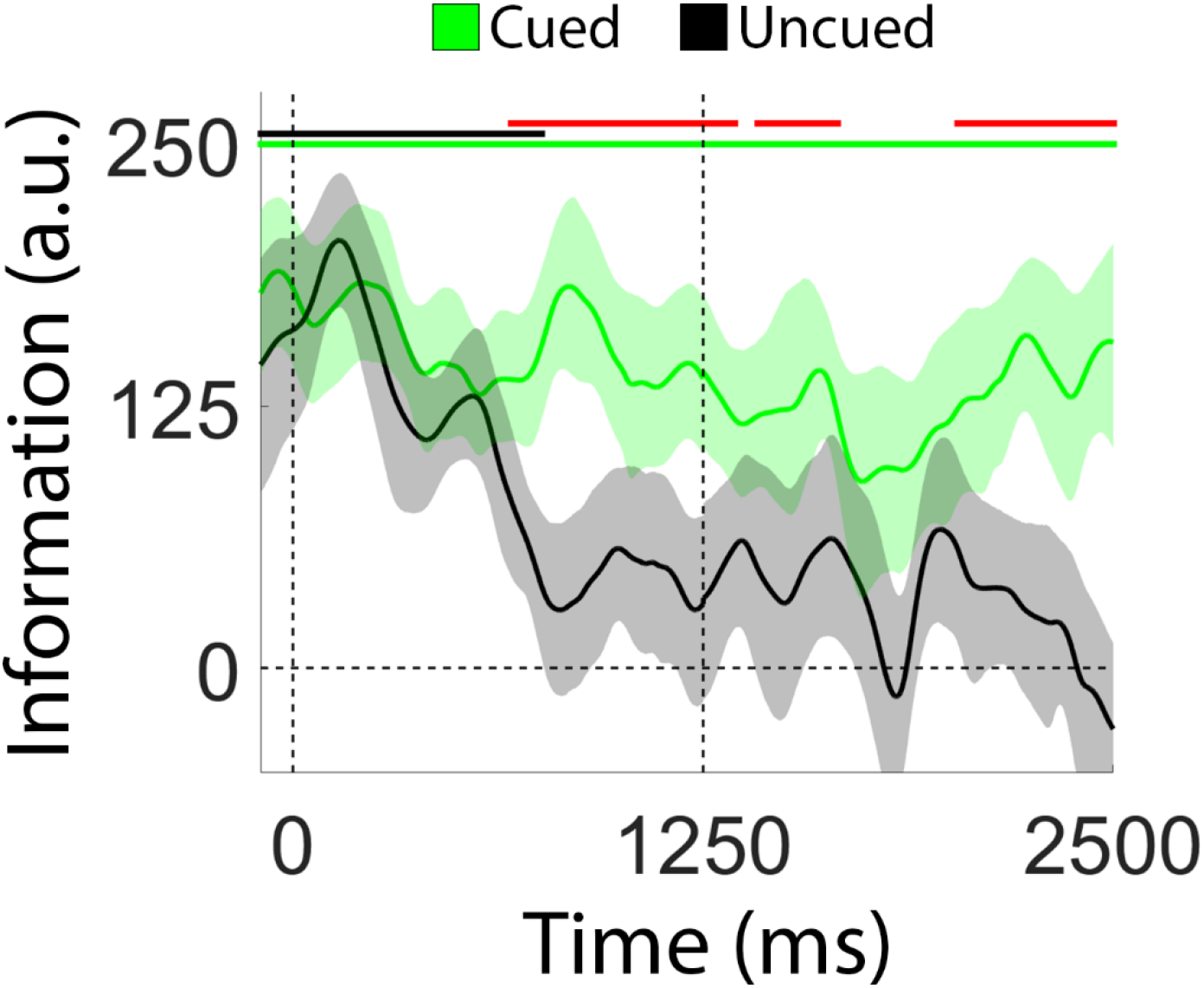
Information Loss is Prevented by a Retrospective Cue Presented Immediately After Encoding. During VE trials, a 100% reliable retrospective cue indicated which disc (blue or red) participants would be asked to report. During these trials we observed a rapid decrease in location information for the uncued disc (black line), but no change in location information for the cued disc (green line). Green and black bars at the top of the plot mark epochs where location information was reliably greater than 0 for the cued and uncued discs, respectively. Red bars mark epochs where location information was reliably larger for the cued relative to the uncued disc. Shaded regions depict 95% within-participant confidence intervals. a.u., arbitrary units.

To summarize, we observed a monotonic decrease in location information with time when participants were required to hold two locations in WM (Figure 5). A retrospective cue presented immediately after termination of the sample display eliminated this decline for the cued location and hastened it for the uncued location (Figure 6). The observation that a retrospective cue mitigates information loss for a cued item is consistent with behavioral and physiological findings suggesting that retrospectively cued shifts of attention insulate cued WM representations from subsequent degradation due to interference or decay (e.g., Pertzov et al. 2013).

### Recovery of Location Information Following a Delayed Retrospective Cue

In a recent study, Sprague et al. (2016; see also Rose et al. 2016; Wolff et al. 2017) documented an apparent recovery of location-specific information following a delayed retro-cue. We tested for a similar effect by examining the effect of a delayed retrocue on location information (Figure 7). During VL trials a neutral cue instructed participants to remember the locations of both discs. Halfway through the delay period the neutral cue changed colors to either blue or red, indicating with 100% probability which disc would be probed. As shown in Figure 7A, location information decreased gradually over the course of the first delay period (0-1250 ms after the sample display; linear slope = −120.78 units/sec; *p* < 0.002, bootstrap test) and continued to decline for the uncued disc during the second delay period (black line). Conversely, location information for the cued item appeared to increase during the second delay period (green line). To evaluate these changes, we computed average location information for the cued and uncued disc after dividing the second delay period into early and late epochs (1250-1700 ms and 2050-2500 ms after sample offset, respectively; Figure 7B) based on visual inspection of the plots shown in Figure 7A. Estimates of location information were identical for the cued and uncued discs during the early epoch and diverged during the late epoch. Direct comparisons of information estimates during the early and late epochs revealed reliably larger information estimates for the cued disc (*p* < 0.007) and reliably lower information estimates for the uncued disc (*p* < 0.015) during the late epoch. We examined sources of location information recovery via a curve fitting analysis. Specifically, we computed and quantified the amplitude and concentration of time-averaged reconstructions of the cued disc during the early and late portions of the second delay period (i.e., 1250-1700 ms and 2050-2500 ms; Figure 7C). Consistent with earlier findings (e.g., Sprague et al. 2016) we observed significantly larger amplitude estimates during the late relative to the earlier epoch (Figure 7D; *p* < 0.001; the difference between concentration estimates during the early and late periods was not significant; p = 0.0543). Thus, in addition to protecting representations of cued items from subsequent degradation (Figure 5), under some circumstances a retrospectively cued shift of attention can directly enhance the representation of a cued item (Figure 7).

**Figure 7.**
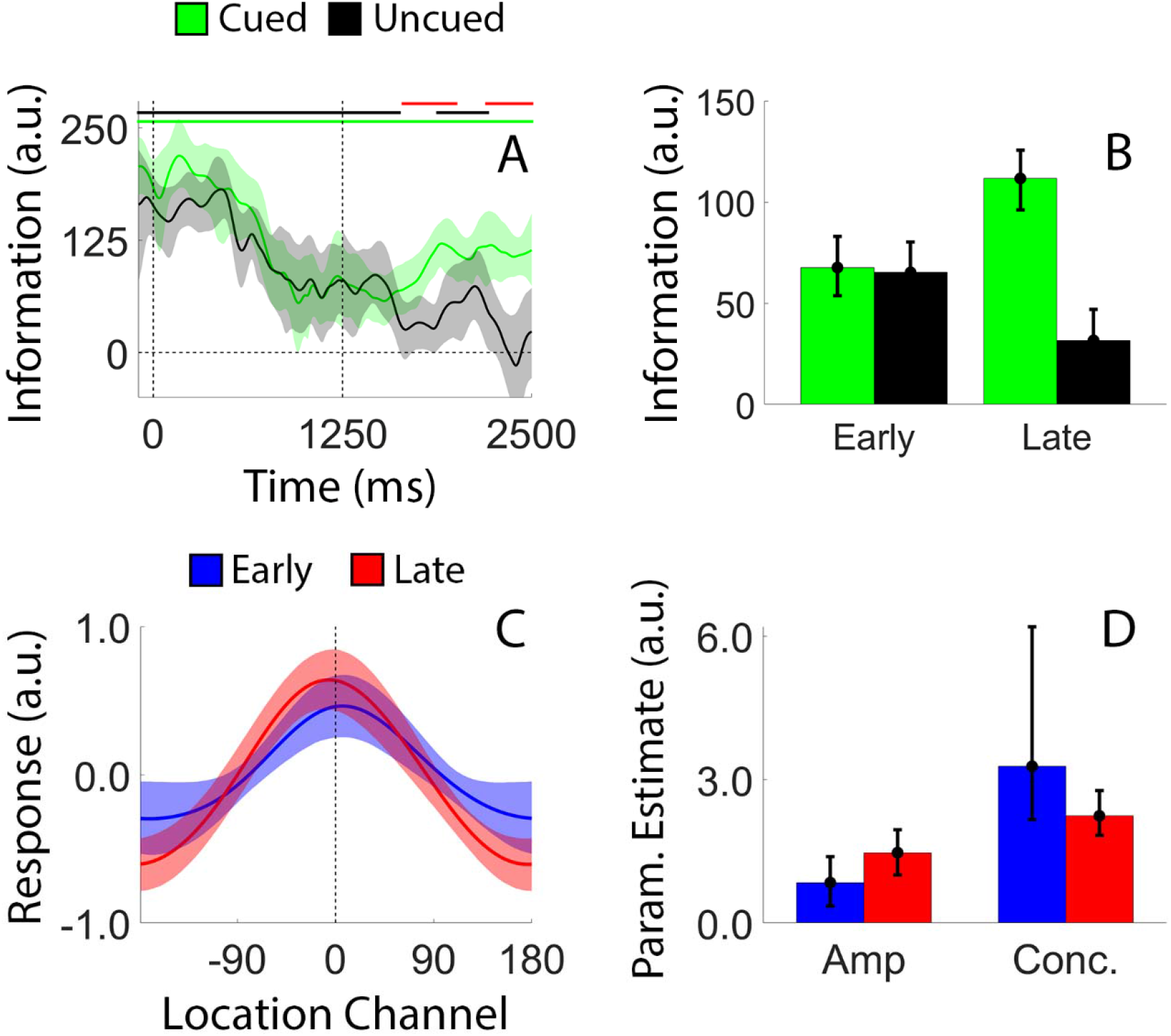
Recovery of Location Information Following a Delayed Retrospective Cue. During VL trials, a neutral cue was replaced by a valid cue midway through the delay period. (A) We observed a monotonic decrease in location information during the neutral portion of the trial, followed by a partial recovery of location information for the cued disc during the valid portion of the trial. Pink, green, and black lines at the top of the plot mark epochs where estimates of location information were reliably greater than 0, while red lines mark epochs where estimates of location information were reliably greater for the cued relative to the uncued disc. (B) To quantify changes in location information after retrocue onset we divided the second delay period into separate early and late epochs (1251-1700 ms and 2050-2500 ms after sample offset, respectively) and computed the average location information for each disc across both epochs. Location information was significantly greater for the cued disc during the late relative to the early epoch, while location information was significantly smaller for the cued disc during the late relative to the early epoch. (C) We computed and quantified time-averaged reconstructions of the cued disc’s location during the early and late epochs of the second delay period. (D) Reconstruction amplitudes were reliably larger during the late relative to the early epoch, consistent with an attention-based enhancement of the cued representation. For all plots shaded regions and error bars depict 95% within-participant confidence intervals.

### Ruling Out Contributions from Eye Movements

We identified and excluded trials contaminated by horizontal eye movements greater than 2.5° (based on a normative voltage threshold of 16 μV/°; Lins et al. 1993). Nevertheless, small but consistent bases in eye position may have contributed to reconstructions of stimulus location. We examined this possibility in two complementary analyses (see *Eye Movement Control Analyses*, Methods). First, we computed angular estimates of eye position from horizontal and vertical EOG recordings and constructed circular histograms of eye position as a function of remembered stimulus location (Figure 8A). Distributions of eye position were remarkably similar across stimulus locations, suggesting that the location-specific reconstructions shown in Figures 3-7 cannot be solely explained by subtle biases in eye position. This conclusion was further supported by a complementary analysis in which we regressed time-resolved estimates of horizontal EOG activity onto remembered stimulus locations (Figure 8B). Time-resolved regression coefficients were indistinguishable from zero across each trial, again suggesting that systematic biases in eye position are insufficient to account for the location-specific reconstructions reported here.

**Figure 8.**
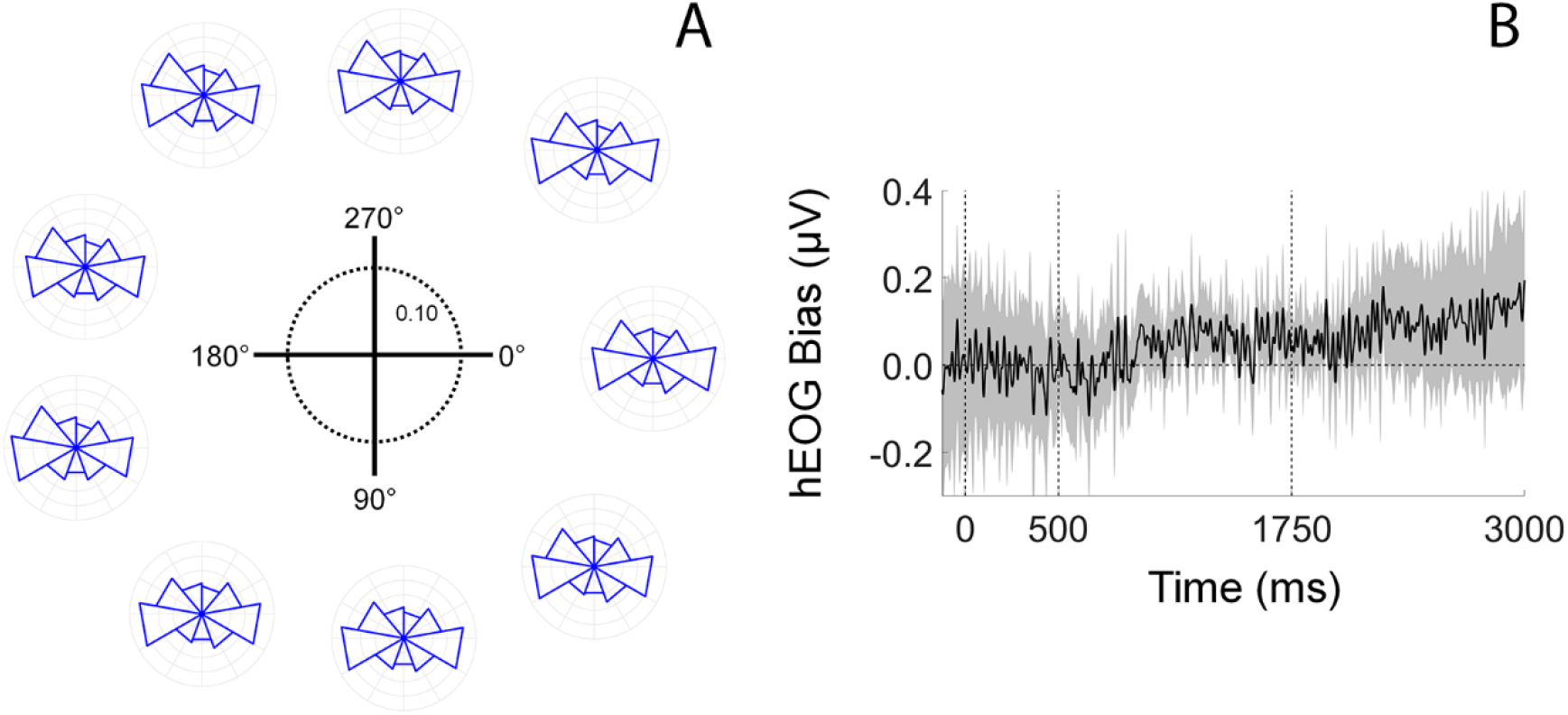
Location-specific Reconstructions cannot be Explained by Subtle Bases in Eye Position. Our analyses focused on VE trials as this is where systematic biases in eye position should be most apparent (i.e., because participants were only required to remember one location). (A) Circular histograms showing angular estimates of eye position are plotted as a function of stimulus location. Each histogram corresponds to one of the 9 possible stimulus locations (see Figure 3A). Data are scaled according to the schematic in the center of the plot. (B) Regression coefficients describing the relationship between horizontal EOG voltage (hEOG; μV) and remembered stimulus locations. Vertical dashed lines at 0 and 500 ms mark the onset and offset of the sample display, while the dashed line during at 1750 ms represents the midpoint of the delay period. Shaded regions are 95% within-participant confidence intervals.

## Discussion

WM performance can be improved by a retrospective cue presented after encoding is complete (Griffin & Nobre, 2003; Landman et al. 2003; Souza & Oberauer, 2016; Myers et al. 2017). Several mechanisms have been proposed to explain retrospective cue benefits in WM performance, including the removal of irrelevant information from WM (e.g., Kuo et al. 2012; Souza & Oberauer, 2016), attentional enhancement of the cued representation (e.g., Myers et al., 2015), protection of the cued representation from subsequent decay or interference (e.g., due to competition from other memory representations or subsequent sensory input; Makovski & Jiang, 2007; Pertzov et al., 2013), or retrieval head start (e.g., Souza et al. 2016). Evaluating these alternatives has proven difficult, in part because extant studies examining the effects of retrospective cues have relied almost exclusively on behavioral reports and/or indirect neural signatures of WM storage. In the current study, we overcame this limitation by directly examining changes in spatially-specific WM representations before and after the appearance of a retrospective cue. Our approach was predicated on recent studies demonstrating that topographic distributions of induced alpha-band activity encode precise location-specific information during covert attention tasks (Foster et al., 2017) and spatial WM tasks (Foster et al., 2016), and it allowed us to visualize and quantify cue-driven changes in WM representations with high temporal resolution. Our key findings are summarized in Figure 9. During neutral trials an uninformative retrospective cue instructed participants to remember the locations of two discs across a blank display, and we observed a gradual degradation the strength of location-specific WM representations with time (Figure 4). During VE trials a 100% reliable retrospective cue was presented immediately after termination of the sample display. This cue eliminated time-based degradation for the representation of the cued disc and hastened it for the uncued disc (Figure 5). Finally, during VL trials a neutral retrospective cue was replaced with a 100% reliable retrospective cue midway through the delay period. We observed a gradual degradation in location-specific WM representations during the neutral phase of the trial followed by a partial recovery of location during the valid phase of the trial (Figure 6). Collectively, our findings suggest that retrospective cues can engage multiple mechanisms to minimize and/or reverse information loss during WM storage (Sprague et al., 2015).

**Figure 9.**
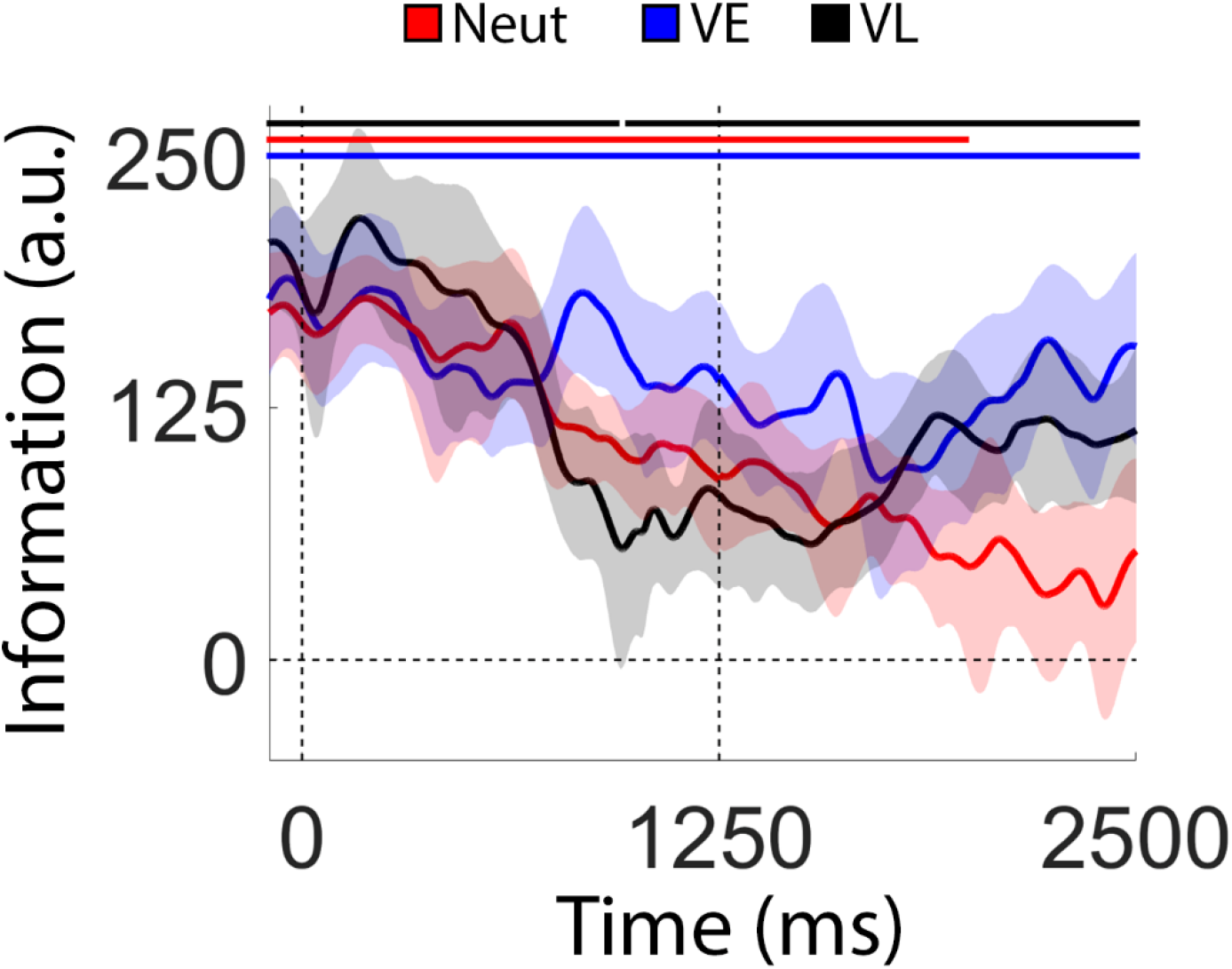
Synopsis of Key Findings. Red, blue, and black lines are reproduced from Figure 5A, Figure 6, and Figure 7A, respectively.

The loss of location information observed during Neutral and VL trials need not imply a loss of memory. Indeed, the fact that participants perform quite well during neutral trials despite an apparent absence of location information suggests at least a partial dissociation between alpha band activity and spatial WM performance. However, there are at least two ways to explain this: one possibility is that spatial WM representations remain stable during the delay period, but location information carried by the alpha-band signal degrades with time. A second possibility is that the memory representations are (partially or wholly) consolidated into a new format not indexed by alpha band activity. The pattern of results we observed during VL trials supports the latter alternative. Under the constraints of information theory (specifically, the data processing inequality theorem; Shannon, 1948), the total information about one variable given the state of another variable (i.e., mutual information) cannot be increased through additional processing. For example, applying multiplicative gain to a noisy spatial WM representation (e.g., by shifting attention to the cued location) would amplify signal and noise to the same extent, resulting in a stronger response but no increase in the information content of the signal (e.g., Sprague et al. 2016; Bays & Taylor et al., 2018). Thus, once location information is lost it cannot be recovered through any amount of additional processing, unless participants have access to an additional source of information. Since there was no external source of location information during the delay period, we can infer that participants were able to access an internal source of information that is not indexed by alpha band activity, including but not limited to an “activity-silent” WM system (Sprague et al. 2016; Rose et al. 2016; Wolff et al. 2017) or long-term memory (Sutterer et al. 2018).

Psychophysical studies suggest that retrospective cues engage mechanisms that insulate or protect cued representations from subsequent degradation (Matsukura et al. 2007; Pertzov et al. 2013). Our findings provide strong support for this view. Specifically, in the absence of a retrospective cue we observed a gradual degradation in location-specific WM representations with time (Figure 4A). However, a 100% valid retrospective cue presented immediately after the encoding display eliminated degradation in the representation of the cued disc and accelerated degradation in the representation of the uncued disc. Rapid degradation of the representation of the uncued disc is nominally consistent with studies suggesting that retrospective cues engage mechanisms that facilitate the removal of uncued items from WM (Astle et al. 2012; Kuo et al., 2012; Williams et al., 2013; Williams & Woodman, 2012). However, it is unclear whether rapid degradation reflects the operation of an active mechanism that purges irrelevant information from WM or the passive (but rapid) decay of information following the withdrawal of attention. This question awaits further scrutiny.

During VL trials, we observed a partial recovery of location information during the second half of the delay period (Figure 6). This result dovetails with several recent empirical studies (Lewis-Peacock et al., 2012; LaRocque et al., 2013; Rose et al. 2016; Sprague et al., 2016; Wolff et al., 2017) documenting a recovery or “resurrection” of decodable stimulus information following a retrospective cue or neurostimulation. Many of these studies have interpreted this recovery as evidence for the existence of an additional, “latent” WM system based on short-term synaptic plasticity that can be used to supplement active storage mechanisms such as sustained spiking activity (e.g., Barak & Tsodyks, 2014; Stokes, 2015). Schneegans and Bays (2017) recently challenged this conclusion by demonstrating that a neural process model based on sustained spiking activity can yield a recovery of location information in a retrospectively cued spatial WM task similar to the one used in this study (see also Sprague et al. 2016). In this model, a colored retrospective cue provides a uniform or homogeneous boost in the activity of one of two neural populations with joint selectivity for a specific color (e.g., red) and location. This boost – coupled with inhibitory interactions between the two neural populations – provides a plausible explanation for the partial recovery of location information following a retrospective cue that was observed during VL trial. However, this model would also predict a boost in location information above and beyond that seen during encoding following the presentation of the retrospective cue during VE trials (see also related empirical findings by Rerko et al., 2014; Souza et al. 2014). Our findings do not support this prediction: we saw no evidence for an increase in location information following presentation of the retrospective cue during VE trials (Figure 5). This implies that there is an upper limit on the information content of reconstructed spatial WM representations that is determined during encoding. Further studies will be needed to explore this possibility in detail.

The absence of a boost in reconstructed spatial WM representations during VE trials (Figure 5) conflicts with psychophysical studies suggesting that retrospectively cued shifts of attention can strengthen WM representations over and above their original encoding strength (e.g., Rerko et al. 2014; Souza et al. 2016). It is also nominally inconsistent with findings reported by Sprague et al. (2016), who observed an increase in the amplitude of a reconstructed spatial WM representation reconstructed from hemodynamic activity when a valid cue was presented immediately after an encoding display (compared to a neutral cue condition). However, the sluggish nature of the human hemodynamic response makes it difficult to infer the source(s) of this effect. Indeed, the data reported by Sprague et al. were acquired with a temporal resolution of 2250 ms, or nearly the length of the entire delay period in the current study. Thus, higher amplitude representations during valid trials could in principle reflect a boost in reconstructed representations over and above their original encoding strength (e.g., Rerko et al., 2014; Souza et al. 2016; Schneegans & Bays 2017) or later degradation in reconstructed representations during neutral trials. Our approach allowed us to disambiguate these possibilities by tracking changes in location-specific reconstructions with a temporal resolution on the order of tens of milliseconds.

Our findings are consistent with studies have documenting links alpha band topography and spatial attention both across and within visual hemifields (e.g., Rihs et al. 2007; Bahramsharif et al. 2010), as well as more recent work demonstrating that momentary changes in alpha band topography can be used to track the locus of spatial attention with high temporal resolution (e.g., Foster et al. 2017). In addition, we have shown that alpha band topography can be used to visualize and track changes in reconstructed WM representations following a retrospective cue. In the absence of a cue we observed a monotonic decrease in memory strength with time. A cue presented immediately after the termination of the encoding display eliminated this decrease and a cue presented midway through the subsequent delay period partially reversed it. Collectively our findings provide new and compelling evidence that depending on circumstances retrospectively cued shifts of attention can (a) prevent subsequent information loss during WM storage, (b) partially reverse prior information loss, and (c) possibly facilitate the removal of irrelevant items from WM.

Author Contributions
E.F.E. conceived and designed the experiment, L.R. and A.N., and E.F.E. collected and analyzed the data. E.F.E. wrote the paper. * - these authors contributed equally to the project and are listed alphabetically.

## Acknowledgements

The authors thank Tommy Sprague for helpful comments on an earlier version of this manuscript.

## References

Astle DE, Summerfield J, Griffin I, Nobre AC. Orienting attention to locations in mental representations. Attn Percept Psychophys 74:146–162 (2012)

Bae G-Y, Luck SJ. Dissociable decoding of spatial attention and working memory from EEG oscillations and sustained potentials. J Neurosci 38:409–422. (2018)

Bahramisharif A, van Gerven M, Heskes T, Jensen O. Covert attention allows for continuous control of brain-computer interfaces. Eur J Neurosci 31: 1501–1508 (2010)

Barak O, Tsodyks M. Working models of working memory. Curr Opin Neurobiol 25:20–24 (2014)

Bays PM, Taylor R. A neural model of retrospective attention in visual working memory. Cognition 100:43–52 (2018)

Blankertz B, Lemm S, Treder M, Gaufe S, Müller K-R. Single-trial analysis and classification of ERP components – A tutorial. NeuroImage 56:814–825 (2011)

Cousineau D. Confidence intervals in within-subject designs: A simpler solution to Loftus and Masson’s method. Tutorials in Quantitative Methods for Psychology. 1:42–45 (2005)

Cowan N. The magical number 4 in short-term memory: A reconsideration of mental storage capacity. Behav Brain Sci 24:87–185 (2000)

Ester EF, Anderson DE, Serences JT, Awh E. A neural measure or precision in visual working memory. J Cogn Neurosci 25:754–761 (2013)

Ester EF, Sprague TC, Serences JT. Parietal and frontal cortex encode stimulus-specific mnemonic representations during visual working memory. Neuron 87:893–905. (2015)

Ester EF, Sutterer DW, Serences JT, Awh E. Feature-selective attentional modulations in human frontoparietal cortex. J Neurosci 36:8188–8199. (2016)

Foster JJ, Sutterer DW, Serences JT, Vogel EK, Awh E. The topography of alpha-band activity tracks the content of spatial working memory. J Neurophysiol 115:168–177 (2016)

Foster JJ, Sutterer DW, Serences JT, Vogel EK, Awh E. Alpha-band oscillations enable spatially and temporally resolved tracking of cover spatial attention. Psychol Sci 28:929–941 (2017)

Griffin IC, Nobre AC. Orienting attention to locations in internal representations. J Cogn Neurosci 15:1176–1194 (2003)

Kleiner M, Brainard D, Pelli D. What’s new in Psychtoolbox-3? Perception (2007)

Kok P, Mostert P, de Lange FP. Prior expectations induce prestimulus sensory tempulates. Proc Natl Acad Sci USA (2017)

Kuo B, Stokes MG, Nobre AC. Attention modulates maintenance of representations in visual short-term memory. J Cogn Neurosci 24:51–60. (2012)

Landman R, Spekreijse H, Lamme VAF. Large capacity storage of integrated objects before change blindness. Vision Research 43:149–164

LaRocque JJ, Lewis-Peacock JA, Drysdale AT, Oberauer K, Postle BR. Decoding attended information in short-term memory: An EEG study. J Cogn Neurosci 25:127–142 (2013).

Lewis-Peacock JA, Drysdale AT, Oberauer K, Postle BR. Neural evidence for a distinction between short-term memory and the focus of attention. J Cogn Neurosci 24:61–79 (2012).

Lins OG, Picton TW, Berg P, Scherg M. Ocular artifacts in EEG and event-related potentials I: Scalp topography. Brain Topography 6:51–63. (1993)

Loftus GR, Masson MEJ. Using confidence intervals in within-subject designs. Psych Bull Rev 1:476–490. (1994)

Luck SJ, Vogel EK. Visual working memory capacity: From psychophysics and neurobiology to individual differences. Trends Cogn Sci 17:391–400 (2013)

Ma WJ, Husain M, Bays PM. Changing concepts of working memory. Nat Neurosci 17:347–356 (2014)

Makovski T, Jiang YV. Distributing versus focusing attention in visual short-term memory. Psychon Bull Rev 14:1072–1078 (2007)

Makovski T, Watson LM, Koutstall W, Jiang YV. Method matters: Systematic effects of testing procedure on visual working memory. J Exp Psychol Learn 36:1466–1479 (2010).

Maris E, Oostenveld R. Nonparametric statistical testing of EEG and MEG data. J Neurosci Methods 164: 177–190 (2007)

Matsukura M, Luck SJ, Vecera SP. Attention effects during visual short-term memory maintenance: Protection or prioritization? Perception & Psychophysics 69:1422–1434 (2007)

Myers NE, Walther L, Wallis G, Stokes MG, Nobre AC. Temporal dynamics of attention during encoding versus maintenance of working memory: Complementary views from event-related potentials and alpha-band oscillations. J Cogn Neurosci 27:492–508. (2015)

Myers NE, Stokes MG, Nobre AC. Prioritizing information during working memory: Beyond sustained internal attention. Trends Cogn Sci 21:449–461 (2017)

Pertzov Y, Bays PM, Joseph S, Husain M. Rapid forgetting prevented by retrospective attention. J Exp Psychol: Hum 39:1224–1231 (2012)

Poch C, Campo P, Barnes GR. Modulation of alpha and gamma oscillations related to retrospectively orienting attention within working memory. Eur J Neurosci 40: 2399–2405. (2014)

Rerko L, Souza AS, Oberauer K. Retro-cue benefits in working memory without sustained focal attention. Memory & Cognition 42:712–728 (2014)

Rihs TA, Christoph MM, Thut G. Mechanisms of selective inhibition in visual spatial attention are indexed by alpha-band EEG synchronization. Eur J Neurosci 25:603–610 (2007)

Rose NS, LaRocque JJ, Rigall AC, Gosseries O, Starrett J, Myering EE, Postle BR. Reactivation of latent working memories with transcranial magnetic stimulation. Science 354:1136–1139 (2016).

Samaha J, Sprague TC, Postle BR. Decoding and reconstructing the focus of spatial attention from the topography of alpha-band oscillations. J Cogn Neurosci 29:1090–1097.

Schneegans S, Bays PM. Restoration of fMRI decodability does not imply latent working memory states. J Cogn. Neurosci 29:1977–1994 (2017)

Shannon CE. A mathematical theory of communication. Bell Syst. Tech. J. 27:379–423 (1948)

Souza AS, Rerko L, Oberauer K. Getting more from visual working memory: Retro-cues enhance retrieval and protect from visual interference. J Exp Psychol Hum 42:890–910. (2016)

Souza AS, Oberauer K. In search of the focus of attention in working memory: 13 years of the retro-cue effect. Atten Percept Psychophys 78:1839–1860. (2016)

Souza AS, Rerko L, Oberauer K. Unloading and reloading working memory: Attending to one item frees capacity. J Exp Psychol Hum 40:1237–1256 (2014)

Sprague TC, Ester EF, Serences JT. Reconstructions of information in visual spatial working memory degrade with memory load. Curr Biol 24:2174–2180 (2014)

Sprague TC, Saproo S, Serences JT. Visual attention mitigates information loss in small- and large-scale neural codes. Trends Cogn Sci 189:215–226. (2015)

Sprague TC, Ester EF, Serences JT. Restoring latent visual working memory representations in human cortex. Neuron 91:694–707 (2016)

Stokes MG. Activity-silent working memory in prefrontal cortex: A dynamic coding perspective. Trends Cogn Sci 19:394–405.

Sutterer DW, Foster JJ, Serences JT, Vogel EK, Awh E. Alpha-band oscillations track the retrieval of precise spatial representations from long-term memory. bioRxiv doi: https://doi.org/10.1101/207860

van Ede F. Mnemonic and attentional roles for states of attenuated alpha oscillations in perceptual working memory: A review. Eur J Neurosci. DOI 10.1111/ejn.13759 (2017)

van Moorselaar D, Foster JJ, Dutterer DW, Theeuwes J, Olivers CNL, Awh E. Spatially selective alpha oscillations reveal moment-by-moment trade-offs between working memory and attention. J Cogn Neurosci 30:256–266 (2018)

Vogel EK, McCollough AW, Machizawa MG. Neural measures reveal individual differences in controlling access to working memory. Nature 438:500–503 (2005)

Williams M, Hong SW, Kang M-S, Carlisle NB, Woodman GF. The benefit of forgetting. Psychon Bull Rev 20:348–355 (2013)

Williams M, Woodman GF. Directed forgetting and directed remembering in visual working memory. J Exp Psychl Learn Mem Cogn 38:1206–1220 (2012)

Wolff MJ, Jochim J, Akyürek EG, Stokes MG. Dynamic hidden states underlying working-memory-guided behavior. Nat Neurosci 20:864–871 (2017).

